# Live Eco-AI, Electrophoresis-Correlative Data-Dependent Acquisition with Artificial Intelligence-Based Data Processing Democratizes Single-Cell Mass Spectrometry Proteomics

**DOI:** 10.1101/2025.04.16.649123

**Authors:** Bowen Shen, Fei Zhou, Peter Nemes

## Abstract

Single-cell mass spectrometry (MS) recently emerged with unprecedented sensitivity to characterize cellular proteomes. Still, limited access to sensitive and fast mass spectrometers impedes its broad adoption, due chiefly to budget constraints. Here, we democratize single-cell MS to a budget-conscious alternative with contemporary performance. A custom-built inexpensive capillary electrophoresis (CE) platform electrophoresis-correlatively (Eco) sorted peptide m/z signals. Detection re-employed a decade-old, slower yet affordable quadrupole-quadrupole-orbitrap (Q-Exactive Plus, QE+, Thermo). Data-dependent acquisition (DDA) with narrow, 1.6-to-4 Th isolation tracked the m/z trends in ∼100% efficiency, outperforming the data-independent reference. The ensuing chimeric spectra were deciphered using artificial intelligence (AI) via CHIMERYS (Thermo). Live Eco-AI augmented our QE+ to ∼15 peptides/spectrum, on par with more advanced analyzers (Lumos and Exploris compared). 1 ng HeLa proteome digest gave 2,142 proteins, topping 969 on a contemporary LC Qq-OT-ion trap reference (Fusion Lumos, Thermo). From ∼250 pg, approximately a cell, 1,799 HeLa-proteins were detected in <15-min-effective electrophoresis. To establish proof of principle, Live Eco-AI was employed to profile 1,524 proteins among differentially fated single stem cells (50-to-75 micrometer diameter) in the *Xenopus laevis* (frog) blastula. The quantitative data revealed previously unknown proteome reorganization during differentiation into the dorsal and ventral lineages.

## INTRODUCTION

Single-cell mass spectrometry (MS) is currently in the limelight of unprecedented research. Ongoing technical efforts aim to deepen the experimentally detectable portion of the proteome by isolating cells, processing their proteomes, and mass-analyzing them in ever-increasing efficiency (reviewed in ^1–2^). Short analytical time frames are desired to improve statistical confidence through analyzing entire cell populations, one cell at a time. Without a technical ability to amplify whole proteomes, access to mass spectrometers with leading sensitivity, resolution, and speed is critical to deep proteome coverage.

Single-cell MS amalgamates various technical advances. Following early work on hemoglobin detection in erythrocytes,^3–4^ capillary electrophoresis (CE) MS pioneered the discovery form of single-cell proteomics in manually dissected large-to-middle, 250–75-µm-diameter embryonic stem cells (blastomeres).^5^ Despite an overwhelming ∼90% yolk protein content in these cells across early development^6–7^, the detectable proteome depth could be deepened by simplifying the sample preparation workflow^8^ and engaging nanoLC-MS^9–10^ to upscale the analyzed proteome amount from the entire cell. Automated cell sorters quickly followed suit, refining single-cell MS to higher numbers of small, ∼10–75-µm cells. Nanoliter-processing systems (e.g., nanoPOTS^11^ and nPOP^12^) down-scaled bottom-up proteomic workflows to sub-microliter volumes. Chemical barcoding elevated the throughput of single-cell proteomics^8^, paving the way to today’s high-throughput single-cell MS using nanoLC^13^. Fast separation in nanoLC^14^, CE^15^, and ion mobility^16–17^ steered analyses to ∼15 min of effective separation, sufficiently fast to entertain single-cell MS for an emerging clinical setting.^18^ Naturally, shorter analytical time frames required increased speed, sensitivity, and resolution, further taxing access to top-of-the-line instruments capable of meeting these technical needs.

Advances in data acquisition and processing offer a partial remedy. In the bottom-up workflow, proteins are measured based on the sequencing of proteotypic peptides, and this process is controlled to ensure reproducibility. Early on, our laboratory^19–21^ and others^22–24^ employed data-dependent acquisition (DDA) to identify up to ∼1,700 proteins among single blastomeres in 30 min in embryos of the South African clawed frog (*Xenopus laevis*).^21, 25^ A DDA ladder ranking ions by abundance partially combated rapid exhaustion in the MS/MS duty cycle, identifying ∼35% more proteins from limited neuronal proteomes of the mouse than the conventional DDA.^26^ By co-fragmenting many peptide features within broad m/z windows, data-independent acquisition (DIA) expanded the empirical single-cell proteome coverage to ∼1,600 proteins from ∼130 μm *X. laevis* blastomeres (∼6 nL cytoplasm)^9^ and up to ∼2,000–3,000 proteins in mammalian cells^17, 27–28^ using new-generation nanoLC-MS. Likewise, DIA enhanced CE-MS to ∼1,100 proteins from single-HeLa-cell-equivalent proteomes and profiled ∼1,200 proteins in ∼15 min from single *X. laevis* blastomeres.^15^ Ion mobility improved protein numbers by 50% in addition.^29^ In prioritized SCoPE, peptide ranking based on identification success, spectral purity, and biological significance improved MS^2^ duty cycle utilization.^30^ Slice-PASEF targeted peptide precursors based on ion mobility, quantifying 1,417 proteins in single HeLa-cell-equivalent proteomes.^31^ Recently, wide window acquisition (WWA) was combined with the artificial intelligence (AI)-assisted CHIMERYS on a new-generation nanoLC-MS system, averaging ∼2,000 proteins per cell.^32–34^ Single-cell MS can now achieve deep proteome coverage, provided a qualifying mass spectrometer is available.

This project aimed to democratize single-cell MS by developing a budget-conscious alternative that matches contemporary proteome coverage. We recognized CE as an ideal platform to separate peptides efficiently, reproducibly, and, importantly for access, considerably cheaply compared to modern instruments. We have found that, unlike complex solvent delivery systems requiring frequent maintenance, electrophoresis is driven by inexpensive and robust power supplies; CE realizes major savings over time vs nanoLC. Following our published protocols,^8, 25, 35^ we set out to build and validate the “affordable” single-cell CE-ESI platform from readily available parts. Further, we recently introduced electrophoresis-correlative (Eco) peptide (m/z signal) sorting, which improved DIA coverage by 38% more proteins, despite still only utilizing <50% of the m/z–MT for identifications.^36^ To reap additional sensitivity from a decade-old but inexpensive Q-q-OT system (Q Exactive Plus, QE+, Thermo), we proposed to address the current inefficiency in MS^2^ sampling in CE-MS by leveraging predictable m/z evolution via Eco-sorting. As with any new technology, protein detection sensitivity and quantitative reproducibility needed testing, configuration, and validation using an authentic, validated proteome standard. To gauge the platform’s potential impact for biology research, we leveraged live Eco-AI to test cellular proteome remodeling during differentiation to form the neural tissue vs epidermal-fated lineages in the chordate *X. laevis* embryo.

## RESULTS

Our technical goal was to deepen single-cell proteome coverage on an affordable budget. Our experimental strategy is overviewed in **Figure 1**. Complementing nanoLC (reviewed in ^1, 37–38^), we strategically selected capillary zone electrophoresis (CE) to separate proteome digests. Our CE-ESI platform, custom-built and validated as per details elsewhere^8, 25^, ensured detection with ultrahigh sensitivity and reproducibility while significantly saving costs during acquisition, operation, and maintenance vs the nanoLC reference. Additionally, we re-employed a more than a decade-old Q-q-OT mass spectrometer (Q-Exactive Plus, **Methods**) to obtain a “high” but slow spectral resolution (70,000 FWHM) at a fraction of the cost compared to the modern instruments. We leveraged Eco-sorting^36, 39^ to simplify MS control by returning to data-dependent acquisition (DDA), a decades-old proven technology on most mass spectrometers. We employed an AI-driven data analysis software, CHIMERYS (Thermo, **Methods**) to extract peptide signals from the resulting highly chimeric tandem mass (MS^2^) spectra. Eco-AI, the combined method, was to be tested on the HeLa proteome digest, with technology validation against a recent LC Q-OT-q-ion trap architecture (Orbitrap Fusion Lumos, **Methods**). We sought to demonstrate Eco-AI by exploring the reorganization of the cellular proteome as the dorsal-animal midline (called D11) and ventral-animal midline (called V11) blastomeres gain neural vs epidermal fates during blastulation of the *X. laevis* embryo.

**Figure 1.**
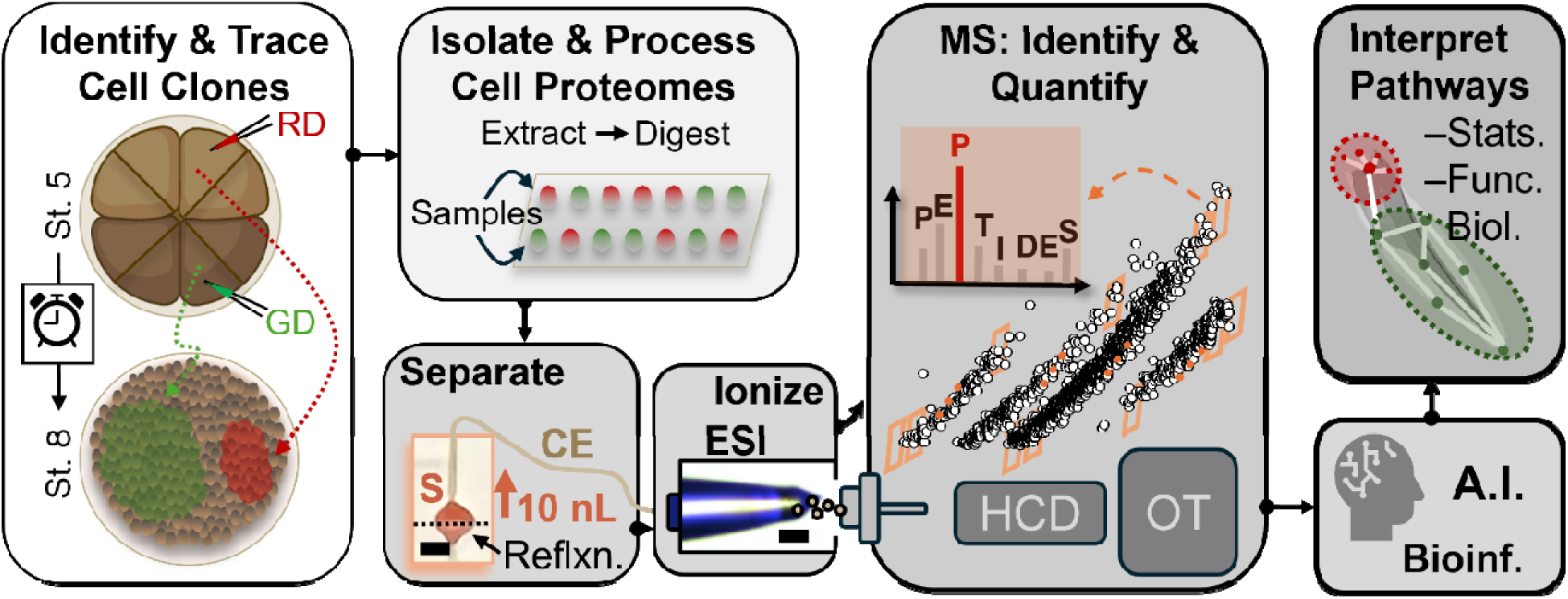
Our experimental strategy establishes affordable single-cell proteomics through capillary electrophoresis (CE), electrospray ionization (ESI), and mass spectrometry (MS). To test proof of principle, the neural tissue fated dorsal-animal midline (D11) and epidermis-destined ventral-animal midline (V11) blastomeres were labeled in red/green fluorescent dyes (RD/GD) in the 16-cell *Xenopus laevis* (frog) embryo. Their descendant lineages were isolated from the blastula (Nieuwkoop-Faber stage 8), phenotyped for the fluorescence markers, and processed for proteome-profiling. Electrophoresis-correlative (Eco) ion sorting was tailored to a decade-old quadrupole-quadrupole-orbitrap (OT) mass spectrometer (Q Exactive Plus, Thermo) for sequencing through higher-energy collisional dissociation (HCD) under real-time (live) control using data-dependent acquisition (DDA). The resulting highly chimeric spectra were deciphered via artificial intelligence (AI)-guided spectral processing (CHIMERYS, Thermo). Live Eco-AI matched the sensitivity of a modern tribrid mass spectrometer (nanoLC on the Fusion Lumos, Thermo). As proof of test, the D11 and V11 descendant cells were profiled to explore proteome reorganization in stem cell populations during differentiation to dissimilar tissue fates. (Created with BioRender.com).

### Strategic Design Considerations for Eco-AI

Our first objective was to chart the rules of successful single-cell proteomics in CE-MS on our chosen “archaic” mass spectrometer against the modern nanoLC tribrid reference. **Figure 2** pursues the mechanistic differences. Preliminary DIA experiments using CE-ESI with the QE+ identified 1,578 proteins from ∼10 ng of HeLa proteome vs 1,357 proteins from ∼10 ng on the nanoLC-ESI Lumos. In agreement with our recent finding,^29^ the peptides from the HeLa proteome digest were Eco-sorted into charge-dependent trends over m/z–MT (**Fig. 2A**), but negligibly so in reversed-phase nanoLC-ESI (**Fig. 2B**). During experimentation, we selected all charge states for sequencing to deepen the experimental proteome coverage. These data are shown and analyzed in **Figure 1** in the **Supporting Information** document (**Fig. S1**). With the +2-charge state accounting for >80% of the empirical proteome identification, we can limit the discussion to this charge state in this report for simplicity. Peptide separation was remarkably different via electrophoresis than retention chromatography.

**Figure 2.**
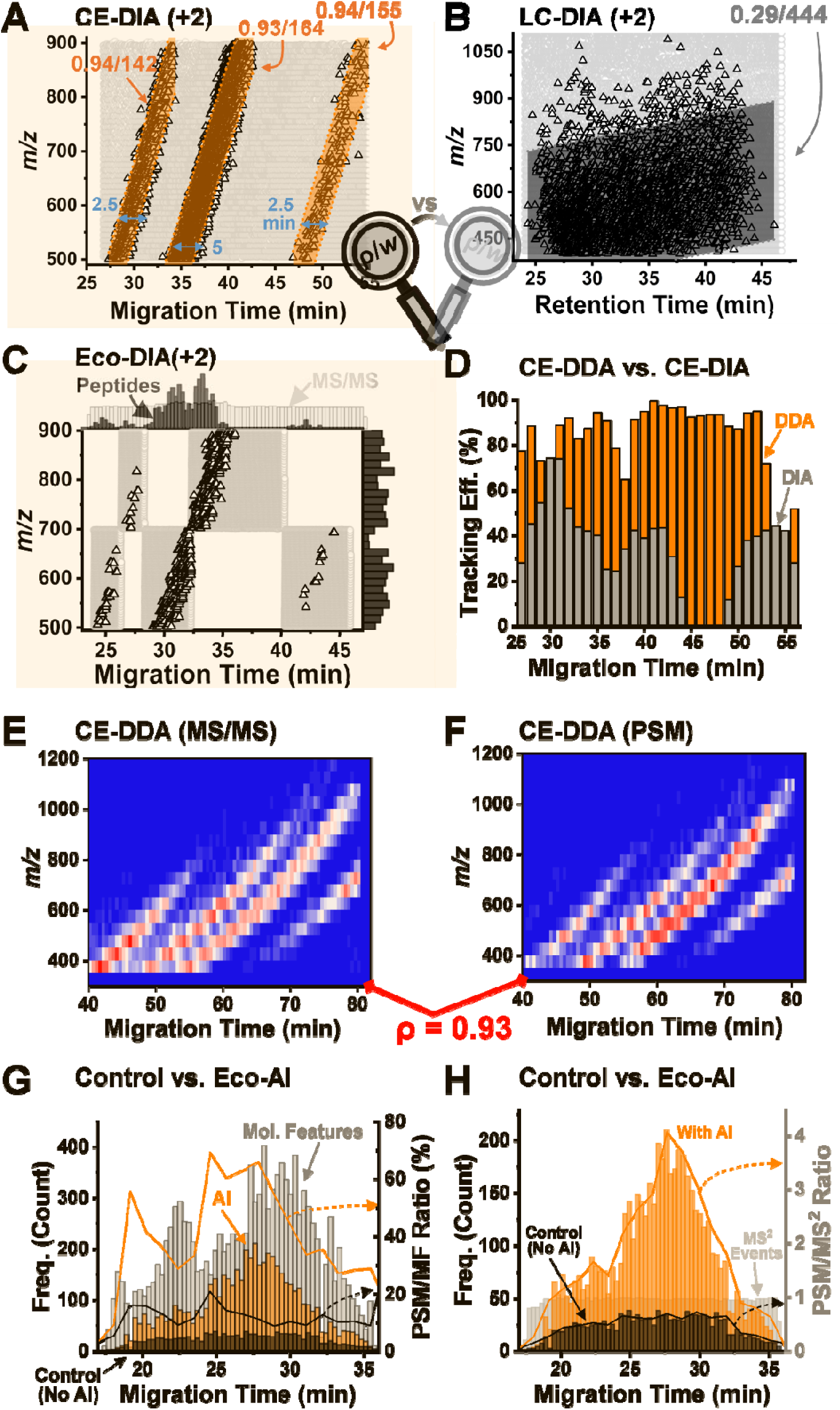
In pursuit of mechanistic idiosyncrasies toward deeper proteomics. 10 ng of the HeLa proteome digest was analyzed using CE and nanoLC MS. While **(A)** electrophoresis-correlative (Eco) sorting ordered the peptide ions by m/z into three correlated temporal clusters (Pearson correlation, ρ ≥ 0.93), **(B)** nanoLC scattered them broadly (ρ = 0.29). **(C)** Eco-DIA segmenting the m/z–migration time space with 5 analytical windows returned ∼38% more proteins than the control, albeit utilizing <50% of the analytical bandwidth. **(D)** DDA using a default m/z window (1.6 Th shown) tracked the m/z–migration time correlations, with efficiencies reaching up to 99% success. **(E)** The emerging MS^2^ events in DDA dynamically aligned with **(F)** the density of the identified peptides (ρ = 0.93), confirming high sequencing success in real time (live). **(G)** AI-assisted data processing (CHIMERYS tested) boosted the success of identifying molecular features (MF) into peptide spectrum matches (PSMs), despite transient peptide ion fluxes yielding highly chimeric tandem mass spectra (MS^2^). **(H)** AI-assisted data processing extracted more than tripled the extractable peptide information from the same chimeric dataset. These enhanced performance metrics suggested deeper proteome depths feasible via live Eco-AI.

Since separation performance de facto limits proteome coverage, Eco-sorting was likely to improve consequences. **Figure 2** gauges this impact between CE (**Fig. 2A**) and nano-LC (**Fig. 2B**). Eco-ordering to have shepherded these precursor ions into three m/z–MT flocks of highly correlated peptide signals, each closely tailing the other (Pearson product-moment, ρ > 0.93). Intriguingly, >95% of these peptide ions migrated along 3, rather narrow trails (ρ ≥ 0.93 each), with values widths (*w*) below 142 Th (∼27–34 min), 164 Th (∼34–43 min), and 155 Th (∼46–55 min). In contrast, the peptide m/z values were dispersed indiscriminately over time in nanoLC, with >95% encompassed within a 444-Th-wide m/z field (**Fig. 2B**, ρ ∼ 0.29). This behavior in CE-MS essentially segmented the m/z–MT landscape into domains of no (grey background formed by the data) and high (PSMs in orange highlight) utility to identify the peptide spectral matches (PSMs) from tandem MS. We recognized the latter as a path to fast sequencing toward deep proteomics and the former as a sacrifice from it. Such patterns were negligible during LC-DIA, where the m/z signals distributed broadly over retention time. These data guided our notion that data acquisition modalities optimized to stochastic m/z–RT signal evolution from nanoLC measurements do not necessarily address high transient fluxes of nominal isobaric signals emerging during CE; the contemporary strategies are not designed to handle Eco-sorting.

We sought to exploit this unique feature of CE separation to tune up the MS sequencing efficiency. We recently deepened proteome coverage by sampling the MS^2^ events along the observed m/z vs. MT trend lines, as closely as possible, ideally in real time (“live”).^36^ As a proxy, we adapted 5 analytical Eco-DIA windows to sample the m/z–MT flocks, as marked in **Figure 2C**. The strategy improved the MS^2^ utilization by ∼50%, quantifying 846 protein groups from 1 ng of the HeLa proteome digest. However, with Eco-DIA, less than 50% of the sampled m/z–MT area contained peptide signals; most of the analytical bandwidth was wasted.

For maximal performance, the Eco trends should be traced dynamically for maximal efficiency, capturing the m/z values with a sufficient but not too large window. Linear regression from **Figure 2A** found an ∼1 Th/s increase in the migrating peptide m/z values. Our recent study revealed peptide coelution within a 2–4-Th wide m/z window.^40^ We recognized the age-old DDA method to be able to sample these trends live, forming the method of Eco-DDA-AI, or “live Eco-AI” for the sake of brevity. The success of DDA depends on the width of the peptide precursor m/z signal isolation; the other key factors, including MS^1^-abundance-thresholding of the MS^2^ transitions and the collision energy were set to the default in this study. **Figure 2D** gauges the efficiency of the commanded m/z signal actually tracking, overlapping with the measured m/z– MT trend. 100% efficiency would mean the ideal condition of all MS^2^ scans sampling peptide signals, whereas 0% efficiency would mean a “complete miss.” The 10-wide m/z-isolation captured the signals in less than half the time (∼45% efficiency) in DIA. In stark contrast, the narrow 1.6-wide m/z windows using DDA, the default setting, were followed with up to 99% efficiency. Further, molecular features (MFs) were dynamically sampled in the m/z–MT dimensions (**Fig. 2E**), yielding PSMs (**Fig. 2F**) in high correlation (ρ = 0.93). These results revealed highly efficient sampling of the Eco-sorted peptides for sequencing.

The quality of the sequencing of MS^2^ data was another question. **Figure 2G** surveys the migrating MFs and their assignment to PSMs. Only 10%–20% of the available peptide-like MFs (see **Method**) were identified as peptides. SEQUEST, a popular and validated search algorithm, extracted barely 1 peptide per MS^2^ spectrum. As CE Eco-ordered peptides into narrow m/z trends (recall **Fig. 2A**), isobaric co-isolation during MS^2^ was expected to yield chimeric spectra.

Recent advances in artificial intelligence (AI) provided a means to extract multiple peptide signatures from the chimeric spectra. Indeed, the CHIMERYS algorithm^41^ found up to 15 PSMs per MS^2^ spectrum in our experiments. The rudimentary, 60-s-wide data binning in **Figure 2G** found the data extraction rate tracking the emerging MFs dynamically, and real time. The elevated sampling rate quadrupled the efficiency of identifying a PSM. On average, approximately three times more peptides were identified per MS^2^ spectrum from the same DDA file processed using AI than without (**Fig. 2H**). AI was a natural fit for our highly chimeric CE-MS data.

### Performance Validation For Single-Cell Proteomics

In the first approximation, the empirical proteome coverage was controlled by the quality of the signals in the MS^2^ spectra. **Figure 3A** evaluates this relationship by refining the precursor m/z isolation window and the OT resolution, while keeping identical cycle times. With 1 ng of the proteome, estimating to ∼4–5 single cells, narrower quadrupole isolation windows (*w*) improved protein identifications. Elevated spectral resolution at lower duty cycle (lower topN) benefited peptide identifications. At the optimum, 1,483 ± 136 proteins were measured on average in 15 min of effective separation, or 2,142 proteins among the triplicates. The identified proteins are tabulated in **Table 1** in the **Supplementary Information** document (**Table S1**). For ∼250 pg of the proteome, approximating to a cell, a slightly wider isolation window (4 Th) proved ideal (**Fig. 3A**). These improvements resulted from higher peptide information content captured in the chimeric MS^2^ spectra using the wider *w* condition (**Fig. 3B**). Up to 15 peptides were identifiable from a spectrum using the 4-Th isolation window. However, too broad m/z isolation (8 Th tested) diminished returns due to increasing spectral interferences (**Fig. 3C**, *p* < 0.001). The optimal *w* in our CE-MS study was broader than the classical ∼0.8–1.6 Th employed in classical DDA, yet narrower than recent efforts employing 15–25 Th widths in DIA^15, 42–44^ and 8–12 Th in WWA DDA^32–33^ in nanoLC. These differences in mechanisms and optimal conditions underscore the importance of tailoring data acquisition to CE-MS.

**Figure 3.**
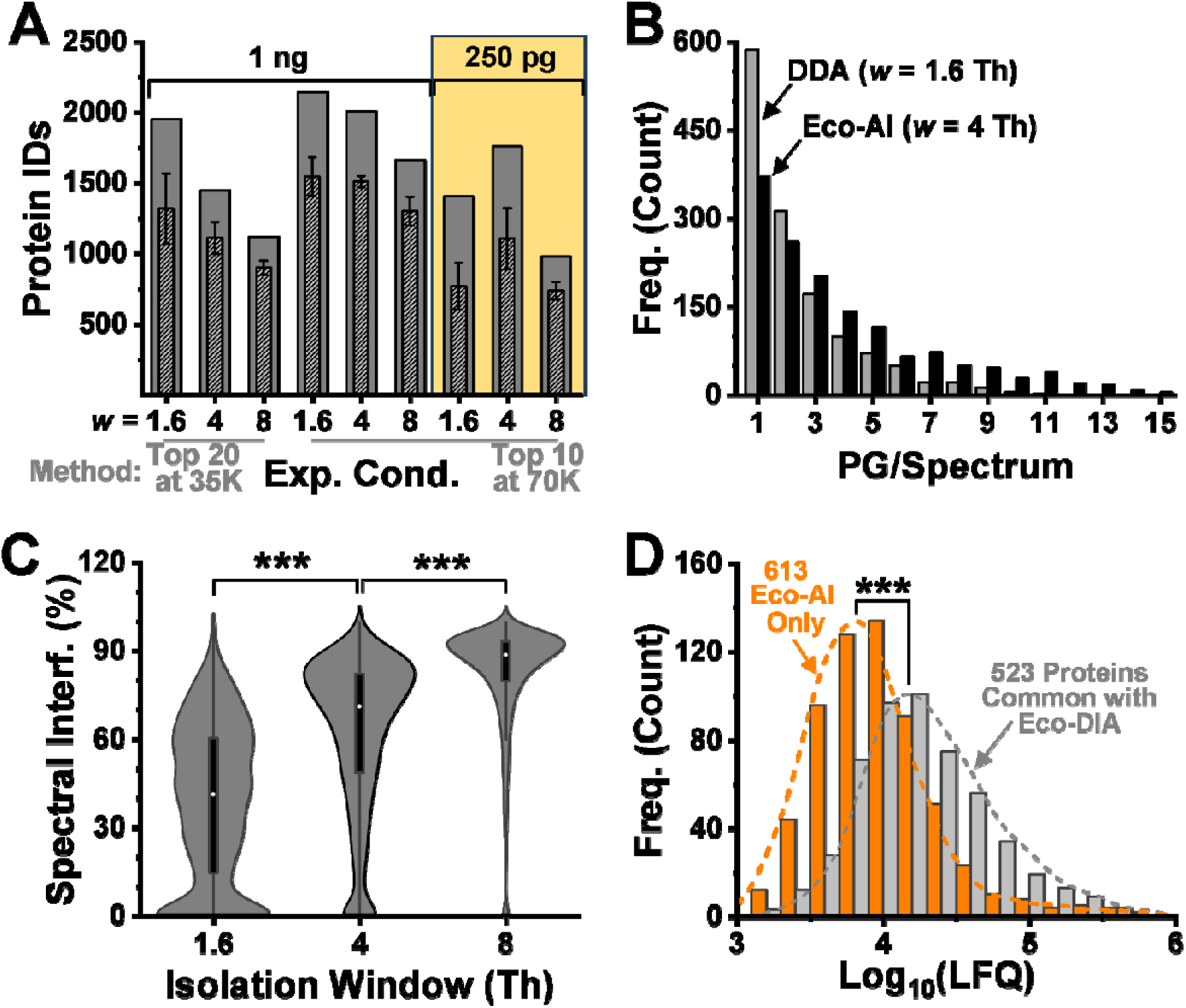
Configuration of the empirical proteome depth. 1 ng and 250 pg of the HeLa proteome digest were analyzed with Eco-AI at different experimental settings at an identical MS cycle duration. **(A)** The quadrupole window of precursor ion isolation (*w*), target ion density (Top 10 vs. 20 ions), and orbitrap spectral resolution (35,000 vs. 70,000 FWHM) were iteratively refined. Eco-AI identified 1,799 proteins in 15 min from ∼250 pg of proteome digest, approximating a single HeLa cell. Quality assessment of the resulting tandem mass (MS^2^) spectra, revealing **(B)** deepening peptide group (PG) information and increasing **(C)** spectra interferences with the wider *w*. **(D)** Label-free quantification (LFQ) confirmed sensitivity enhancement via live Eco-AI vs the CE-DIA reference. Key: ***, *p* < 0.001 (Mann-Whitney *U* test).

The performance of protein quantification was an important metric for biological applications. The classical DDA^21^, DIA^15^, and our last-generation Eco-DIA^36^ approach offered the closest reference technologies for benchmarking. The replicates averaged ∼1,106 ± 217 proteins, or 1,799 proteins cumulatively, without using match-between-runs during data analysis (**Table S2**). For equal proteome amounts, live Eco-AI returned 4.5-times more proteins than the classical DDA and 2.6-fold than the DIA. Compared to the Eco-DIA, live Eco-AI also doubled the number of quantified proteins (**Figure 3D**). 613 proteins that were only quantifiable using Eco-AI dominated the lower dynamic range of the concentration (*p* = 1.54 × 10^−65^) based on the calculated label-free index values used as a proxy^5^. A high Pearson correlation (ρ = 0.92) between the LFQ concentrations measured in 1 ng and 10 ng proteome digests confirmed high quantitative reproducibility (**Fig. S2**). The median coefficient of variation (CV) was ∼16.4% among n = 3 technical replicates after LFQ normalization to the total peptide signal in each sample. These results confirmed that Eco-AI improved proteome coverage (sensitivity) while maintaining good quantitative reproducibility.

The ultimate test was through a match against a more recent nanoLC mass spectrometer available in our laboratory. We selected the Dionex 3000 nanoLC and Fusion Lumos mass spectrometer as the reference. This tribrid system had the upper hand with higher analyzer resolution (70 K vs 120 K FWHM), double sequencing speed (3 Hz vs 7.5 Hz), and improved ion transmission efficiency (sensitivity). At an optimal m/z isolation of 8 Th in the quadrupole (**Fig. S3**), nanoLC-Lumos identified 969 proteins from 1 ng of the proteome digest among n = 5 technical replicates (**Table S3**). These proteins were reproducibly quantified from 1 µL sample using the autosampler (17.3% median CV based on LFQ). Using a comparable, ∼20-min effective separation window, CE-MS measured 2,142 proteins among n = 3 technical replicates. Most of the proteins were detected also by CE (**Fig. 4A**). These successes were due to Live Eco-AI augmenting the slower analyzer (165 MS^2^ scans/min vs. 318 on the Lumos) to sequencing rates, <600 PSMs/min, comparable within the nanoLC-Lumos (**Fig. 4B**). These identification rates exceed recent CE-MS work by us and others. Further, the identification rates from live Eco-AI CE-MS were surprisingly comparable to results on recent-generation, faster analyzer systems employing 2–7-times longer chromatography separations^14, 18, 45^ (**Fig. S4**). Consequently, substantially more peptides can be identified per MS^2^ spectrum with Eco-AI over LC-MS (WWA) (**Fig. 4C**). Put simply, Live Eco-AI polished an old mass spectrometer to single-cell proteomics with matching performance to a new one.

**Figure 4.**
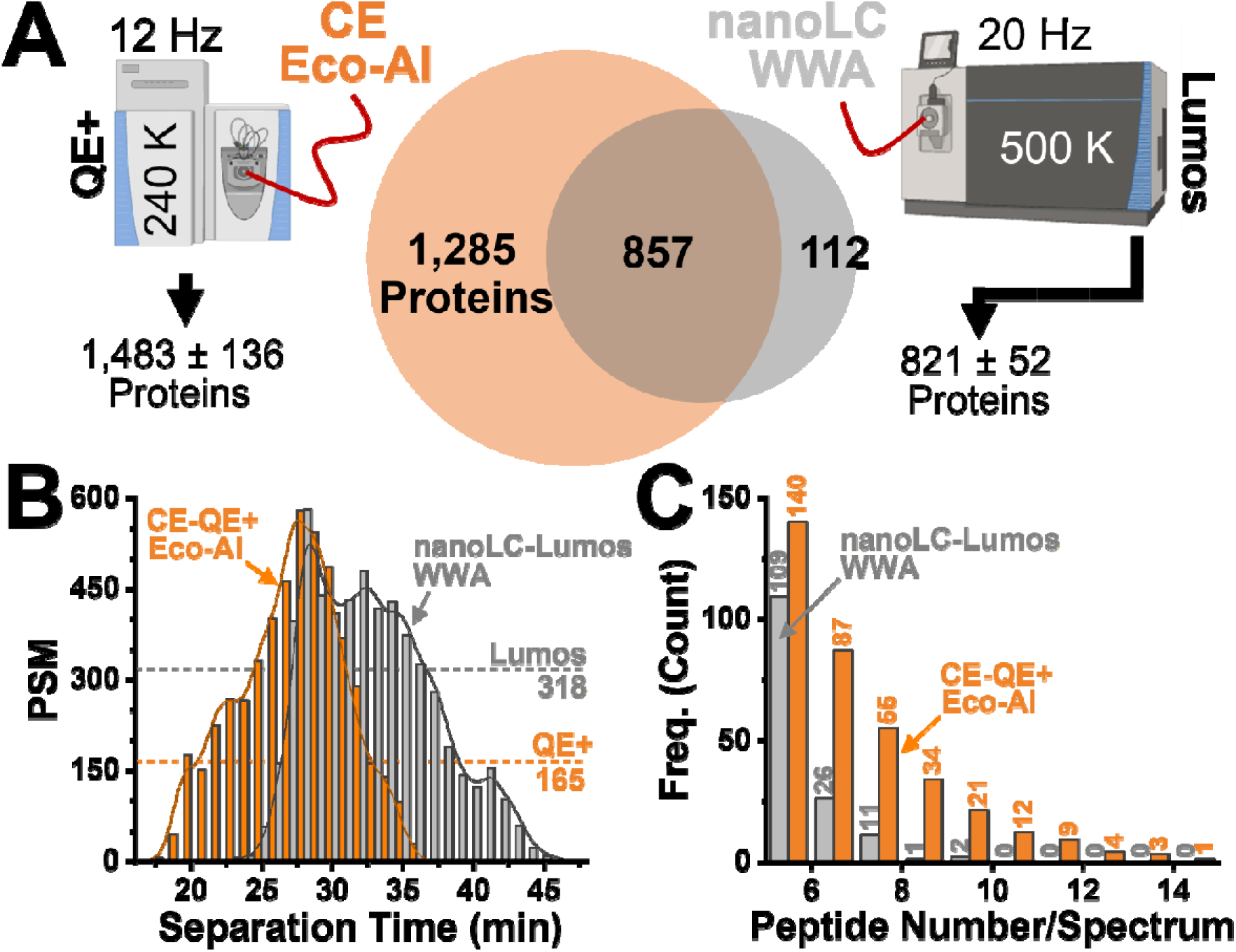
Benchmarking of Live Eco-AI against the nanoLC-tribrid-WWA reference. 1 ng of the HeLa proteome digest was analyzed with CE on the Q Exactive Plus (QE+) and a nanoLC quadrupole-orbitrap-ion-trap tribrid reference (Fusion Lumos, Thermo). **(A)** The comparison of proteome coverage revealed significant sensitivity enhancement via Live Eco-AI on the older and slower mass spectrometer. **(B)** Live Eco-AI boosted the peptide spectral match (PSM) identification rate to be on par with the modern tribrid instrument. **(C)** This improvement was due to deeper peptide information captured in the tandem mass spectra acquired during Live Eco-AI than the nanoLC WWA reference.

### Single-Cell Differentiation in the Embryo

We leveraged Live Eco-AI to profile the proteome of dorsal vs ventral cells in the blastula of the *X. laevis*^46^ embryo (recall **Fig. 1**). **Figure 5A** exemplifies fluorescent microinjection of the D11 (red) vs V11 (green) blastomeres in the 16-cell (stage 5) embryo to label their offspring lineage. These descendant cells respectively give rise to the neural and epidermal tissues in the larva, as demonstrated in our cell-fate tracing experiment in the stage-32 larva. The ∼250-µm-diameter giant precursor blastomeres shrank to ∼50–75-µm diameter by the mid-blastula (stage 8), which are estimated to yield ∼2.5 ng yolk-free proteome in 150 pL of total cytoplasm, on top of an ∼90% of yolk background^6, 47^. After dissociating the embryo following an established protocol^48^, we used a micropipette under a microscope to fluorescently identify and then transfer randomly n = 8 different D11 and n = 8 different V11 descendant cells onto a microscope slide. To aid results interpretation, each cell was obtained from the same clutch laid by the same father and mother, reducing biological variability. Representative cells are imaged in **Figure 5B**. To approximate sensitivity to mammalian somatic cells, only ∼50 pg yolk-free proteome, viz. (∼2%) of the available cell proteome was measured with Eco-AI in this project.

**Figure 5.**
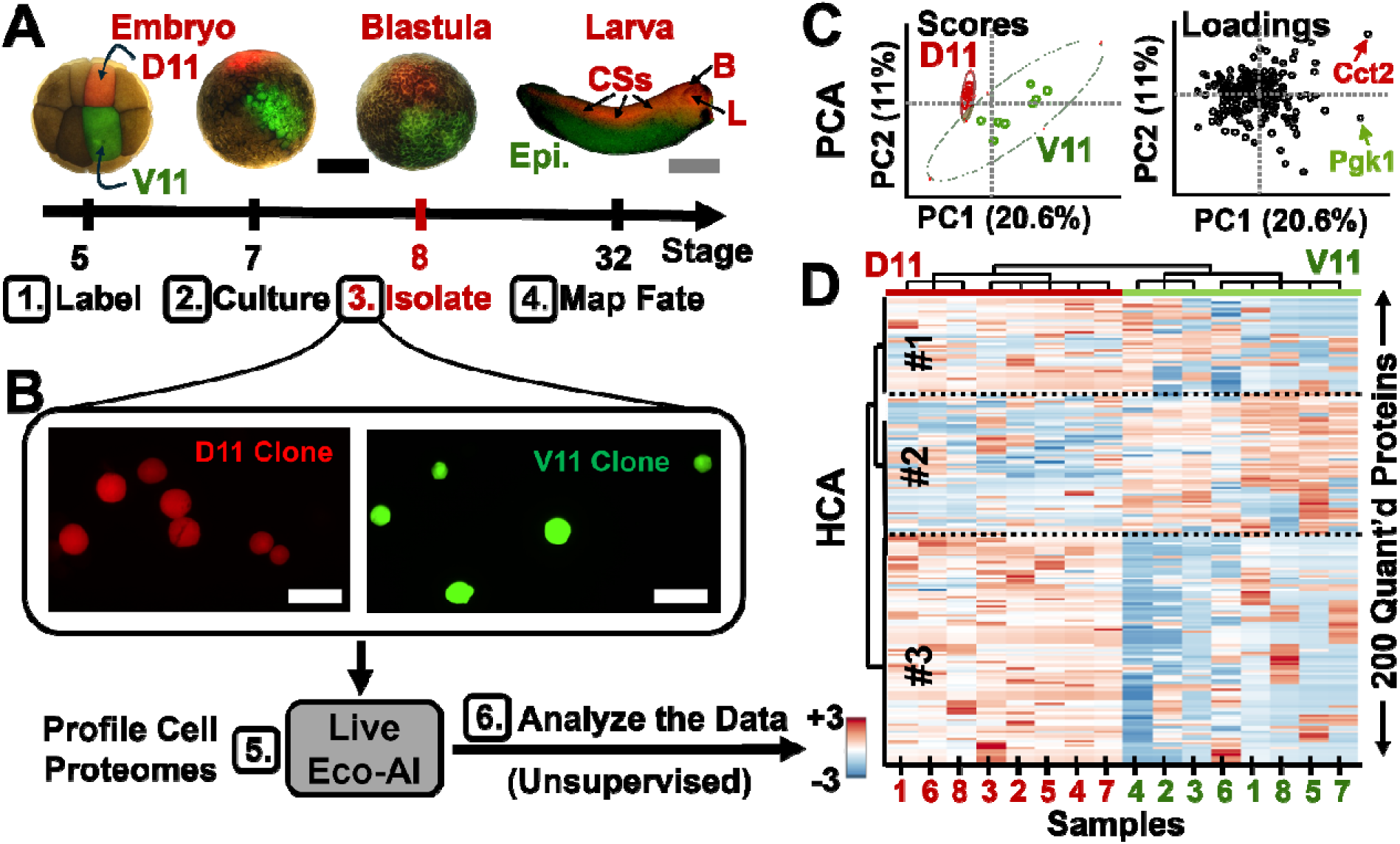
Proteome profiling of single cells differentiating in the *X. laevis* embryo. **(A)** Example for labeling the D11 blastomere in red and the V11 in green fluorescence via microinjection in the 16-cell embryo. These cells established the larva’s neural (CS, central somites) vs epidermal (Epi.) tissues, respectively. **(B)** Representative isolation of the descendant cells (∼50–75 µm in diameter) in the blastula using a micropipette and a fluorescent microscope. 500 pg, ∼2% of the cellular proteome digests were analyzed via Eco-AI. **(C)** Unsupervised multivariate and statistical comparison of the cell proteomes. Principal component (PC) analysis (PCA) revealed systematic differences among the sample types (scores plot), driven by varying protein concentrations (loadings plot). The first 2 PCs are shown (see the text). **(D)** Hierarchical cluster analysis (HCA, z scale shown) confirmed the significance of these differences and clustered the top 200 proteins into groups with different concentration profiles (#1–3). Scale bars, 500 µm (black), 2 mm (gray), 100 µm (white).

The data served as a quantitative palette of the cellular molecular state, potentially in appreciation of functional differences between the dorsal-ventral phenotypes. A total of 1,524 proteins were identified (**Table S4)**. This marked an ∼10-fold sensitivity enhancement at 2–4-fold higher throughput vs custom-built CE platforms, which reported ∼800–1,700 proteins from ∼5–10 ng proteomes on average.^15, 19, 21, 25, 49^ The Live Eco-AI identifications denote at least ∼80-times sensitivity enhancement vs recent nanoLC work, which reported 644–1,650 proteins from 40 ng–2 µg of starting proteomes.^9, 50^ According the PantherDB 18.0^51^, >1,200 of these proteins were annotated to binding (456 proteins), catalytic (489), structural (107), and transporter activities (34). They are also dynamic respondents in metabolism (606), stimuli (112), and cellular localization (128). The data analysis is available in **Figure S5**. **Figure S6** compares the protein LFQ concentrations to the precursor blastomeres using DIA on the same CE-MS platform^15^, revealing significant sensitivity enhancement materialized through Live Eco-AI (p = 5.6 × 10^−15^).

The cells were compared based on their proteome profiles. **Figure 5C** presents the multivariate and statistical analyses of the median-normalized log_10_-transformed LFQ concentrations. In an unsupervised approach, we employed principal component (PC) analysis (PCA) to gauge the samples (scores plot) and their protein amounts (loadings plot). The top PCs described 20.6%, 11%, and 10% of variance in the dataset; we selected the first two for further discussion. The single-cell samples formed two clusters in the PCA scores plot. Upon revealing the identity of the cells, we learned that these groups corresponded to the D11- and V11-derived clones in the blastula. While the neural tissue fated blastomeres formed a tighter cluster, those with epidermal destiny appeared more variable. During early development of the embryo, the descendant D11 lineage undergoes convergent extension to establish neural tissues (retina, brain, central somite), whereas the V11 offshoots scatter to coat the larva (**Fig. 5A**). The PCA loadings plot measured the contribution of the proteins explaining the observed differences. Proteins with comparable abundance among the cell tyles populated the origin. In contrast, those in the periphery were expressed variably. For example, Cct2 (PC1 = 0.25, PC2 = 0.21, *p* = 0.01) were more abundant in the D11 clones, whereas Pgk1 (PC1 = 0.24, PC2 = –0.08, *p* = 0.01) was enriched in the offsprings of the V11.

Significance was confirmed using statistical models. **Figure 5D** shows the hierarchical cluster analysis (HCA) based on the 200 most differential proteins. A close-up of the figure is provided in **Figure S7**. Orthogonal to the PCA results (**Fig. 5C**), this unsupervised method also revealed dissimilar sample types. The two groups corresponded to the D11 vs V11 lineages, as we learned on revealing the identity of the samples at this stage of the data analysis. Based on their abundance profiles, the model clustered the proteins into 3 primary groups (labeled #1–3). The proteins in #1 and #3 were enriched in the dorsal lineage, whereas those in the #2 were accumulated in the ventral. For example, Cndp2 (cluster #1) and Cct2 (cluster #1) were highly enriched in the D11 clones, while Pgk1 (cluster #2) and Grhpr.2 (cluster #2) was more abundant in the V11 clones. We also found the protein enrichment revealed in this study matched previous findings from our lab on the 16-cell embryo^8^: Rpl31 (cluster #1), Calm1 (cluster #1), and Rpl29 (cluster #3) were enriched in the D11 cell clones, whereas Ckb (cluster #2) possessed higher abundance in the V11. As these data may indicate conservation of protein expression across the lineage, we queried the expression of the corresponding genes via Xenbase^52^ as the search engine. Cct3 and Eif5a were concentrated in the neural-tissue-fated dorsal lineage of the embryo. In agreement, *in situ hybridization* found *cct3* upregulated in the neural tube, central nervous system, brain, and eye of the tadpole^53^, and Eif5a is a potent effector of neuron growth.^54^ Conversely, alpha globin larval-5 (Hba-l5) populated the V11 offsprings. Indeed, *hba-15* is accumulated in the ventral blood island^55^, the later developing non-neural ectoderm and axial mesoderm,^56^ and plays a role in the formation of the pronephros^46^, a tissue fate of the V11 cell^46^.The exact functional roles of these proteins are unknown to us. Nonetheless, the data resulting from Live Eco-AI can now be used to appreciate the molecular machinery underpinning cell differentiation more deeply.

Gene Annotation of the identified proteins helped estimate the technology’s potential utility. We mapped the identified proteins to canonical knowledge using PANTHER database 18.0 (**Fig. S5**). Many proteins in all 3 clusters are known components in protein synthesis. They included more than 30 ribosomal proteins and translation initiation factors (e.g., Eif4ai, Eef1b, Eef2, etc.) and chaperone proteins (Cct2, Cct6a) for folding and assembling newly synthesized proteins. The protein functions were enriched in energy metabolism. For example, Cox7a2 and Ndufv1 (both in group #2) are involved in electron transport in mitochondria. ATP synthase subunits (e.g., Atp5f1a, Atp5f1c, Atp5pb, Atp5pd, and Atp5po), enriched in cluster #3, produce ATP in mitochondria. Vdac3 plays an important role in mitochondrial transport. Pgk1 and Gapdh (group #2) are involved in glycolysis, while malate dehydrogenase proteins (e.g., Mdh1 and Mdh2), enriched in group #3, catalyze the conversion of malate to oxaloacetate, an essential pathway in the TCA cycle and energy currency in cell division.

## CONCLUSION

Here, we developed Live Eco-AI, a budget-conscious alternative to single-cell proteomics that matches contemporary performance. We custom-built a CE platform to Eco-sort peptide ions by m/z and developed Live Eco-AI to boost the sequencing efficiency of an old QE+ to the nanoLC Lumos tribrid reference. With efficient cycle time utilization, Live Eco-AI on this high-resolution, slow but affordable mass spectrometer identified 1,799 proteins from a single HeLa-cell-equivalent, ∼250 pg proteome digest in 15-min effective separation. From ∼50 pg of yolk-free proteome, or ∼2% of the *X. laevis* blastomere’s content, Live Eco-AI detected 1,524 proteins in 10 min of effective electrophoresis. The quantitative profiles distinguished cell heterogeneity between the D11 and V11 cell clones at the blastula stage. These results agreed with known proteome asymmetry in the embryo, validating the performance. Despite using an older-generation mass spectrometer, this project obtained the deepest single-cell proteome coverage in a mid-blastula stage *X. laevis* embryo to our knowledge while measuring only a fraction of the cellular contents. With sensitivity enhancement and budget-consciousness, Live Eco-AI democratizes access to deep single-cell proteomics using CE-MS, complementing the strengths of modern nanoLC-MS.

## METHODS

### Materials

HPLC-grade solvents and chemicals, including acetic acid (AcOH), acetonitrile (ACN), formic acid (FA), and methanol (MeOH) were obtained from Thermo Fisher Scientific. Ammonium bicarbonate (AmBic) was purchased from Avantor (Center Valley, PA). The HeLa proteome digest standard (part no. 88329, Pierce, Rockford, IL) and N-dodecyl-β-D-maltoside (DDM, part no. 89903, Rockford, IL) were provided by Thermo Fisher Scientific. The fused silica capillaries were purchased from Polymicro Technologies (40/105 µm inner/outer diameter, part no. 1068150596, Phoenix, AZ) for CE analysis. The CE-nanoESI emitters were fabricated from borosilicate glass capillaries (0.75/1.00 mm inner/outer diameter, part no. B100-75-10, Sutter Instrument, Novato, CA). Proteome digestion was performed with MS-grade Trypsin Platinum (part no. VA900A, Promega, Madison, WI). To minimize analyte loss during sample preparation, all samples were processed in 0.5-mL LoBind vials (Eppendorf, cat no. 022431064, Enfield, CT).

### Solutions

The CE background electrolyte (BGE) was prepared to contain 1 M FA in 25% (v/v) ACN. The CE-ESI sheath solution comprised 0.5% (v/v) AcOH in 10% (v/v) MeOH. The HeLa proteome digest or single cell samples were dissolved in the sample solvent prepared with 0.05% (v/v) FA in 75% (v/v) ACN. The embryo culture media was prepared with 3% Ficoll in 100% Steinberg’s solution. For fluorescent labeling, the fluorescent dextran solution was prepared with 0.5% (v/v) green dextran in DEPC water. For tissue dissociation, the Newport buffer was prepared as described elsewhere.^10, 57^ The proteome lysis solution consisted of 0.2% DDM in 50 mM AmBic.

### Animal Care and Embryology

Sexually mature *X. laevis* frogs were supplied by Xenopus1 (Dexter, MI) or Nasco (Fort Atkinson, WI). All procedures related to the humane care and management of X. laevis were authorized by the Institutional Animal Care and Use Committee at the University of Maryland, College Park (approval no. R-FEB-21-07 or R-FEB-24-05). To capture natural biological variability in the data, the embryos were obtained through the natural mating of one pair of parents. Two-cell embryos presenting stereotypical pigmentation^58^ were cultured to the 16-cell stage (Nieuwkoop-Faber^59^, NF stage 5), where the cell types were readily identifiable based on pigmentation, size, and location in reference to reproducible cell-fate maps.^46^ To trace tissue lineages, the left dorsal-animal midline (termed D11) or the ventral-animal midline (termed V11) cell of the 16-cell embryo was injected with 1 nL (0.5% v/v) of dextran (Alexa Fluor 488, 10,000 g/mol formula weight, anionic, Thermo Fischer). These fluorescent labeled embryos were cultured in 3% Ficoll in 100% Steinberg’s solution at room temperature to the mid blastula stage (NF stage 8).

### Single-Cell Isolation and Proteome Processing

Using sharpened forceps, the vitelline membrane ensheathing the embryo was carefully dissected away to isolate the fluorescence-labeled tissue under a fluorescence stereomicroscope. The biopsy was gently transferred to a glass vial containing Newport buffer, followed by gentle nutation to obtain a cell suspension (24 RPM speed for 10 min, ambient temperature, series no. I2CF61041100, Fisher Scientific). Using a 1 µL pipettor, the cells were carefully transferred onto a culture plate, then swiftly rinsed (in <1 min) with HPLC water to wash off salts and the complex media surrounding the cells. The resulting cells appeared intact based on the emission that the GD label produced in the D11 or V11 cells under a fluorescent stereomicroscope with FITC filter (Nikon SMZ18 with the excitation wavelength at 488 nm). Each cell was lysed in 1 µL of the *proteome lysis buffer* at room temperature for 10 min. The single-cell proteomes were denatured by heating to 60 °C for 15 min, before digestion at 40 °C for 5 h by the addition of 1 µL of 0.1 µg/µL trypsin platinum in 50 mM AmBic. The classical steps of proteome reduction and alkylation were omitted to alleviate sample loss and simplify the workflow. To compensate for liquid evaporation, the Eppendorf vials were kept tightly capped, and the contents of the vial combined via intermittent centrifugation, every ∼15 min. The single-cell samples were dried at room temperature, then stored at –80 °C until analysis.

### CE-ESI-MS Analysis

The single-cell proteomes were analyzed on the same CE-nanoESI platform following the same protocols that we recently described in detail.^15, 20–21^ In this study, ∼1 ng or ∼250 pg of the HeLa proteome digest was electrophoresed at +250 V/cm electrical field strength in a background electrolyte (BGE)-filled 100-cm-long capillary (vs. Earth-grounded capillary outlet). The capillary outlet was connected to an electrokinetically pumped (+500–800 V) sheath-flow CE-nanoESI interface, built following a previous design^60^ and operated in the cone-jet regime^8^ for maximal ionization^61^. A stable Taylor cone was visualized under a long-working-distance objective microscope (Mitutoyo Plan Apo, Edmund Optics, Barrington, NJ) with a CCD camera (EO-2018C, Edmund Optics). This interface was affixed to a 3-axis translation stage to position the tip of the CE-ESI source ∼1 mm from the grounded orifice of a mass spectrometer. The generated peptide ions were measured on a quadrupole–orbitrap tandem high-resolution mass spectrometer (QE+, Thermo Fisher Scientific), under control by data-dependent acquisition (DDA).

Two Top-N methods were devised for CE-MS Eco-AI. In both approaches, survey (MS^1^) scans were obtained with the following parameters: Orbitrap (OT) mass resolution, 70,000 full width at half maximum (FWHM, at m/z 200); scan range, *m/z* 350–1,600; C-trap maximum injection time, 240 ms; AGC target for MS^1^ and MS^2^, 3 × 10^6^ counts (both); charge states triggering MS^2^, +2, +3, and +4. During the “Top-20” method, the 20 most abundant (“top”) precursor ions were fragmented, the fragments C-trapped in <110 ms, and analyzed at 35,000 FHWM resolution in the OT. In contrast, the Top-10 method targeted fewer, specifically 10 “top” precursor ions, thus enabling longer times for C-trapping (<240 ms) and higher-resolution detection (70,000 FHWM). In both data acquisition modalities, the ion signals were dissociated in nitrogen collision gas at 28% normalized collision energy (NCE) in the higher-energy collisional dissociation (HCD) cell. The isolation widths of 1.6, 4, and 8 Th were assessed for each method. The *X. laevis* single-cell proteomes were measured using the Top-10 approach, with a 4 Th isolation window.

### NanoLC-ESI-MS Analysis

The HeLa proteome digest was reconstituted in LC-MS grade water containing 0.1% (v/v) FA. One nanogram of the HeLa proteome digest was trapped (C18 trap column, 100 μm i.d., 5 μm particle with 100 Å pores, 2 cm length, Acclaim PepMap 100, Thermo) at 5 μL/min for 5 min. This proteome digest was separated on a 200 cm µPAC HPLC column (S/N 1100462, Thermo) at a 600 nL/min flow rate. A nanoflow liquid chromatograph (Dionex Ultimate 3000 RSLC, Thermo) provided a 60 min gradient of Buffer B (100% ACN, 0.1% v/v FA) as follows: from 1% to 22.5% over 22.5 min, then from 22.5% to 40% over 7.5 min, then from 40% to 95% over 5 min, and held for 95% for 10 min, then decreased to 1% in 3 min and equilibrated at 1% for 12 min. The peptides were charged in an electrospray source of an electrified (+2,300 V vs. Earth-ground) stainless-steel emitter (part no. ES542, Thermo Fisher). The peptide ions were detected on a quadrupole-ion trap-orbitrap tribrid high-resolution mass spectrometer (Orbitrap Fusion Lumos, Thermo). Peptide signals were surveyed (MS^1^) between *m/z* 380–1,600 at 120,000 FWHM in the Orbitrap analyzer every 3 s. The threshold was set to 2.0 × 10^4^ counts, with AGC normalized up to 250% using auto maximum injection time and a 30 s dynamic exclusion. A DDA method was programmed to isolate the precursor ions within windows of *m/z* 1.6, 4, 8, 12, and 20 Th, followed by fragmentation in the HCD cell at 30% NCE. The resulting MS^2^ spectra were recorded at 60,000 FWHM.

### Data Analysis

The Eco-AI MS data were analyzed in Proteome Discoverer 3.0 (Thermo Fisher Scientific), executing CHIMERYS (Prediction model: inferys_2.1_fragmentation) for protein identification and quantification. The MS^2^ acquisition was extracted from Proteome Discoverer 3.0 (Thermo Fisher Scientific), executing Sequest HT for the conventional DDA analysis without AI. Protein identifications were searched against the HeLa Proteome (UP000005640, downloaded from UniProt in July 2023, containing 20,523 entries). The search parameters included static modification (cysteine carbamidomethylation), dynamic modifications (methionine oxidation), minimum and maximum peptide length (7 and 30 amino acids, resp.), maximal missed cleavage sites (2), fragment ion mass tolerance (20 ppm), and the search for common contaminants (enabled). Similarly, the MS data from experiments on the *X. laevis* were mapped against the *X. laevis* proteome (UP000694892, downloaded in April 2022, containing 42,596 entries), employing the same parameters for the HeLa. Peptide and protein identifications were based on at least 1 proteotypic peptide, filtered to <1% false discovery rate (FDR), computed vs a decoy proteome database enlisting the reversed proteome sequence. For a direct comparison between Eco-AI and DIA performed on CE-MS, protein identification and quantification were obtained with DIA-NN 1.9^62^ with the settings as follows: minimal peptide length, 5; maximal peptide length, 35; maximum missed cleavage, 2; and precursor charge, 2–4. All the other parameters were set to default. The LFQ data were median normalized to compare identical proteome amounts, log_10_-transformed to reduce the quantitative dynamic range with MetaboAnalyst 6.0^63^, and analyzed through statistical means in OriginPro 2020b (Origin Lab, Northampton, MA). The MFs were surveyed in MzMine 3.9^64^ with the settings: m/z range, 380– 1,200; noise level, 10^4^ counts; scans, MS^1^; minimum signal height, 10^4^ counts; *m/z* tolerance, 0.05 Th or 20 ppm. For the non-parametric statistical test, the Mann-Whitney *U* test was performed with R to yield the exact p values unless it reached the limitation.

### Scientific Rigor

Each HeLa proteome digest was analyzed in 3–5 technical replicates (the same sample measured multiple times). A total of n = 16 different blastomeres (biological replicates) were calculated from *X. laevis*. The cell type was identified based on various factors, including pigmentation, size, location, and the fluorescent emission visualized under a stereomicroscope. The biological replicates were analyzed randomly, and the mass spectrometer was calibrated and validated monthly to maintain a good performance.

### Safety

Standard safety protocols were followed when handling all chemicals and biological samples. Careful precautions were taken during the manipulation of capillaries and ESI emitters to mitigate potential puncture hazards. All electrically conductive components of the CE-HRMS platform were grounded or shielded from exposure within an enclosure with a safety interlock.

## Data Availability

All the MS primary files and MS^2^ spectral libraries and the HeLa and *Xenopus* proteomes were deposited in the Proteome Exchange Consortium via the PRIDE partner repository with the data set identifier PXD062702.

## ACKNOWLEDGMENTS

Parts of this research were supported by the Arnold and Mabel Beckman Foundation (Beckman Young Investigator Award to P.N.), the Chan-Zuckerberg Initiative (award to P.N.), or the National Institute of General Medical Sciences (award no. R35GM124755 to P.N.) or the National Institute on Aging (award no. R01AG088147 to P.N.) of the National Institutes of Health.

## AUTHORSHIP CONTRIBUTIONS

P.N. and B.S. designed the study. F.Z. labeled, dissociated, and imaged the single blastomeres. B.S. developed the technology, processed the blastomeres, and measured the blastomere proteomes. B.S. and P.N. analyzed the data and interpreted the results. B.S. prepared the draft report. P.N. revised and finalized the manuscript. P.N. acquired the funding. All the authors commented on the manuscript.

