## Supplementary material for "Live Eco-AI, Electrophoresis-Correlative Data-Dependent Acquisition with Artificial Intelligence-Based Data Processing Democratizes Single-Cell Mass Spectrometry Proteomics": SI Document

### FIGURES

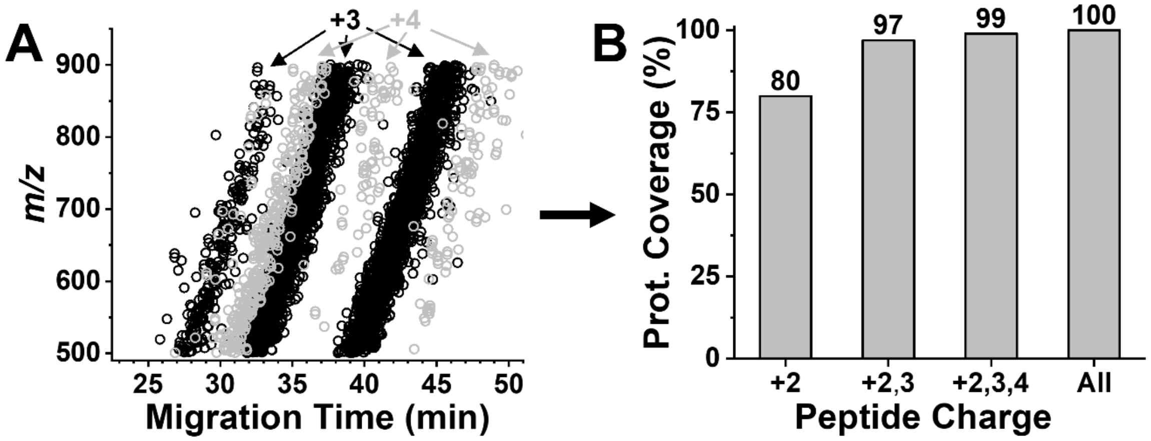

**Figure S1. Peptide separation vs. protein identification. (A)** Eco-sorting at the +3 and +4 charge states (protonated). **(B)** Cumulative coverage of the proteome over all the unknown and +2 or higher charge states measured. Accounting for ~80% of the identified proteome, +2 charge state was selected for results interpretation in this study.

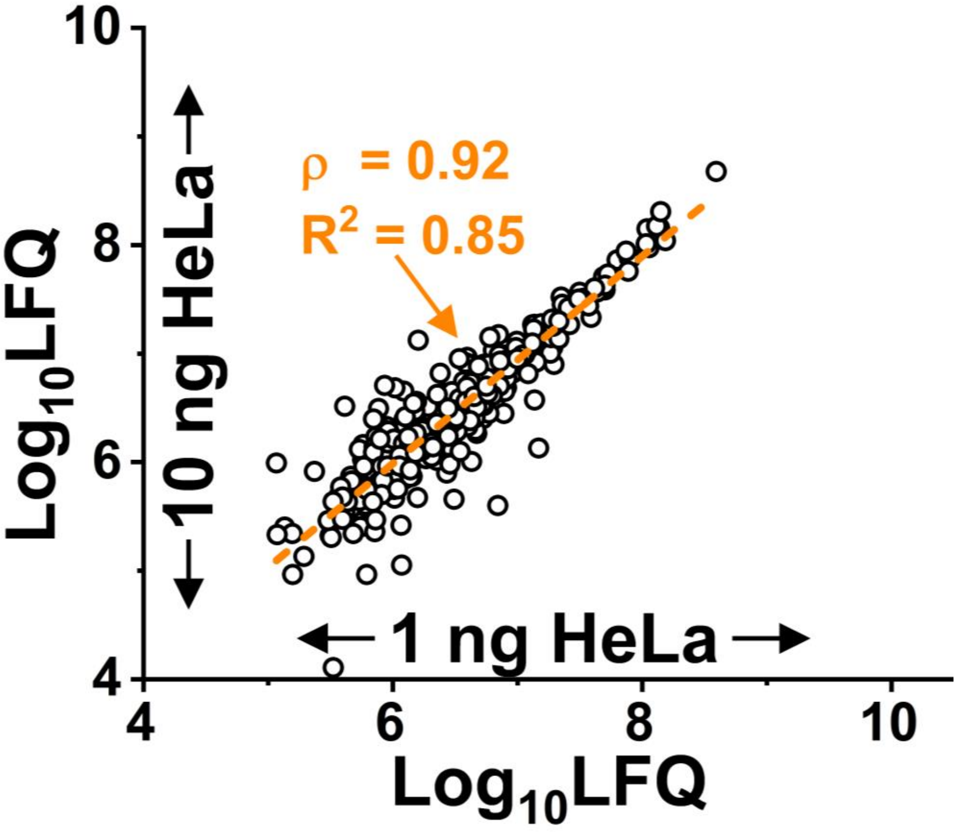

**Figure S2.** Cross-correlation analysis of proteome quantification. 1 ng and 10 ng of the HeLa proteome digest were measured. The protein concentrations were estimated based on LFQ. The high Pearson correlation moments (ρ) and linear regression coefficients ($R^2$) calculated between the LFQ concentrations revealed robust quantification using Live Eco-AI.

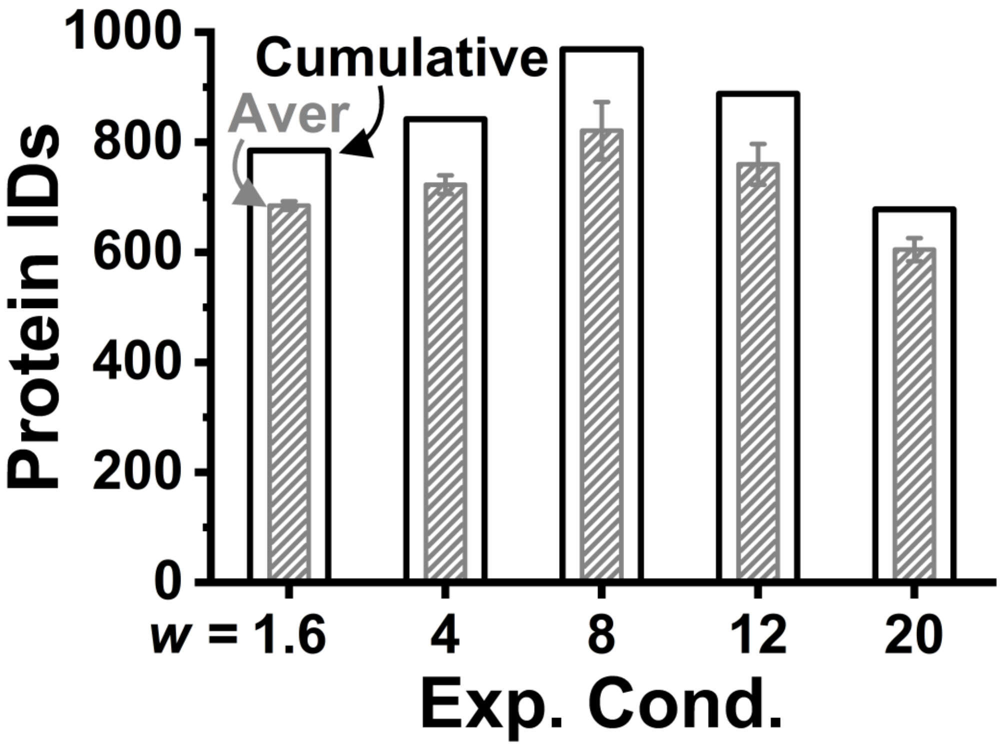

**Figure S3.** Configuration of experimentation conditions (Exp. Cond.) to improve proteome sensitivity using nanoLC-MS. The wide-window acquisition (WWA) method was tested with different quadrupole precursor m/z isolation windows (*w*). 1 ng of the HeLa proteome digest was separated using a 30-min reversed-phase gradient. The separated peptides were ionized in a nano-flow electrospray ionization source and detected on a modern tribrid mass spectrometer (Fusion Lumos, Thermo). Average and cumulative protein identifications are compared among n = 4–5 technical replicates.

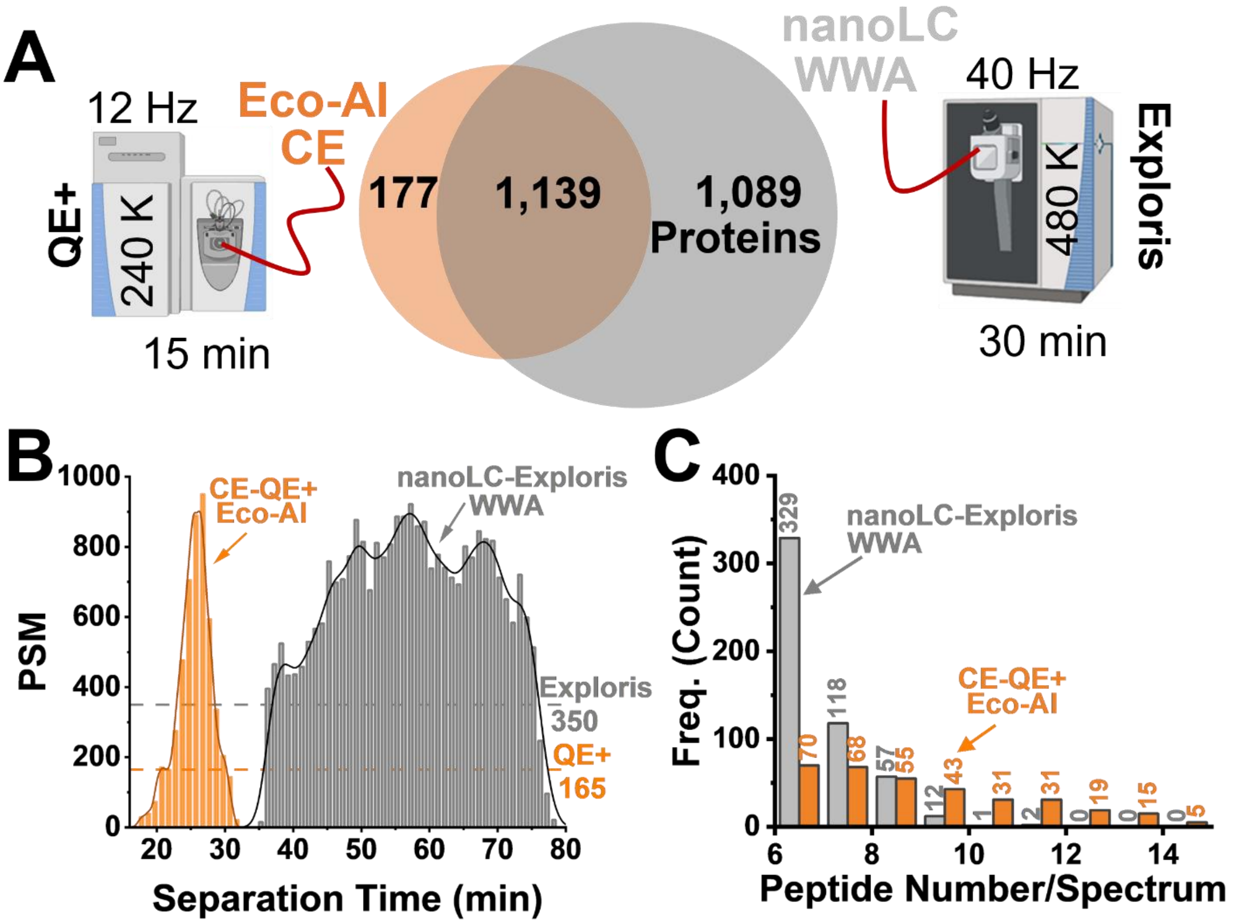

**Figure S4.** Performance benchmarking Live Eco-AI on the Q Exactive Plus (QE+) vs the modern nanoLC Exploris MS reference. Single-cell-equivalent HeLa proteome digests were measured using both platforms (250–200 pg). The nanoLC-MS results were obtained on an Exploris 480 employing wide window acquisition (WWA) in an independent study published elsewhere.[1] **(A)** Although the final proteome depth was higher using the Exploris with elevated sensitivity, resolution, and speed, **(B)** the experimental rate of peptide sequencing was comparably high between CE-MS and nanoLC-MS. **(C)** The Live Eco-AI on the QE+ could extract more peptides per $MS^2$ spectrum despite one-third of the scan rate and the separation time used in the nanoLC-WWA reference. These results demonstrate quasi-matching performance among the technologies.

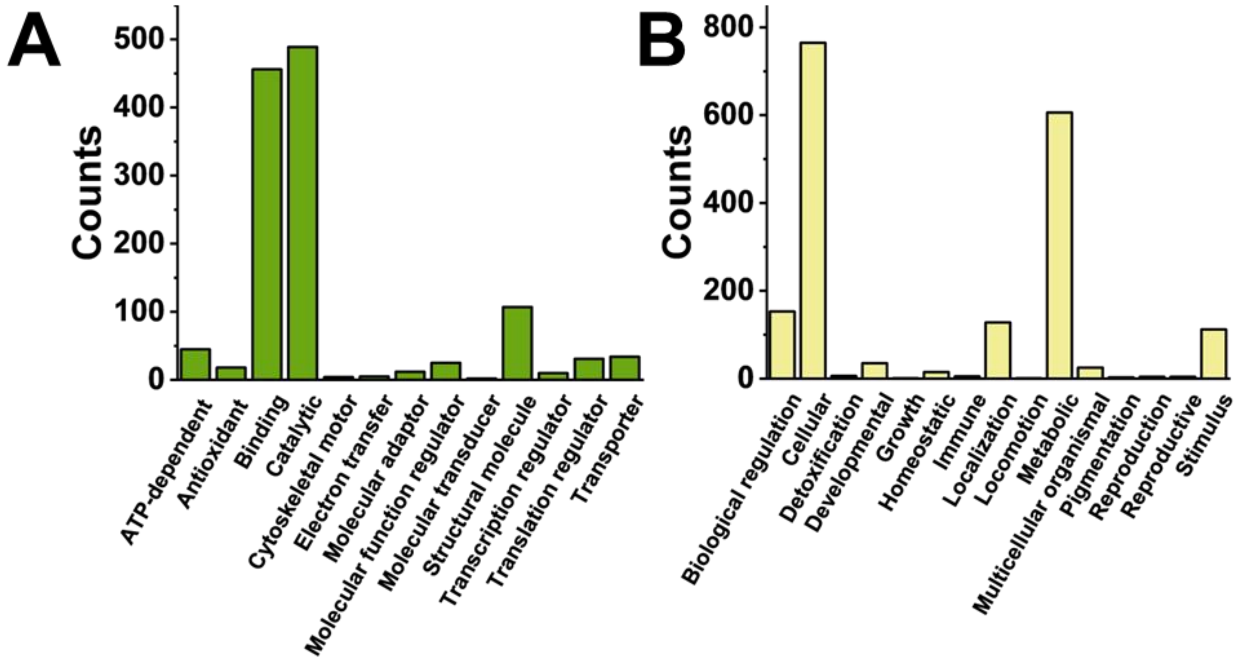

**Figure S5.** Interpretation of canonical knowledge for the proteins identified in the single *X. laevis* blastomeres. The proteins that were identified in ~2% of the single-cell proteome was matched to PantherDB 18.0[2]. Analysis of **(A)** molecular function and **(B)** biological processes among the detected proteins.

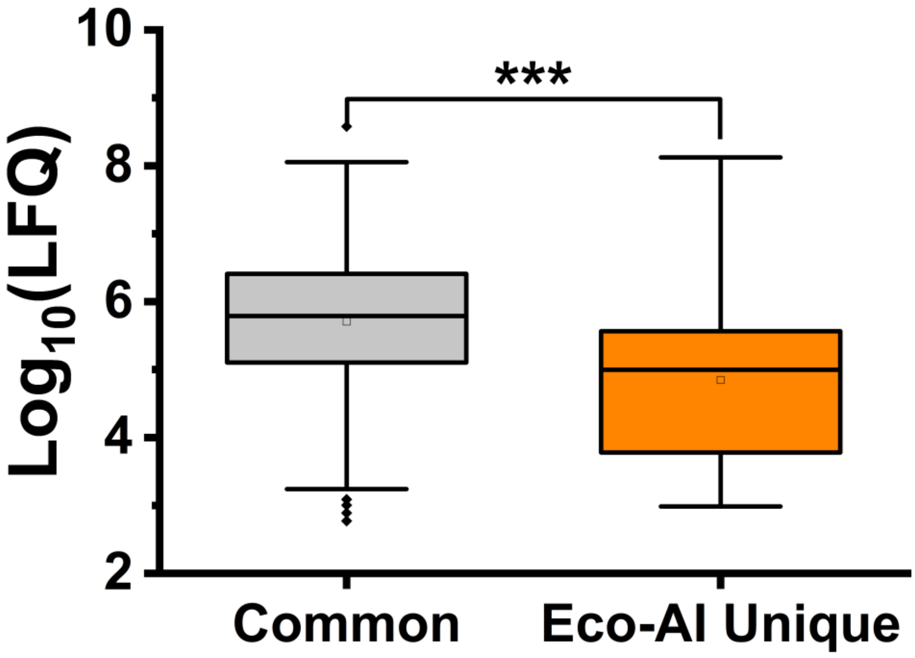

**Figure S6.** Comparison of Live Eco-AI sensitivity against the DIA[3] reference. Live Eco-AI identified proteins in the single *X. laevis* cells that occupied the lower domain of the measured concentration range via label-free quantification (LFQ). These differences were statistically significant, revealing notable sensitivity enhancement via Live Eco-AI in this work. Key: *** $p < 0.001$ (Mann-Whitney *U* test).

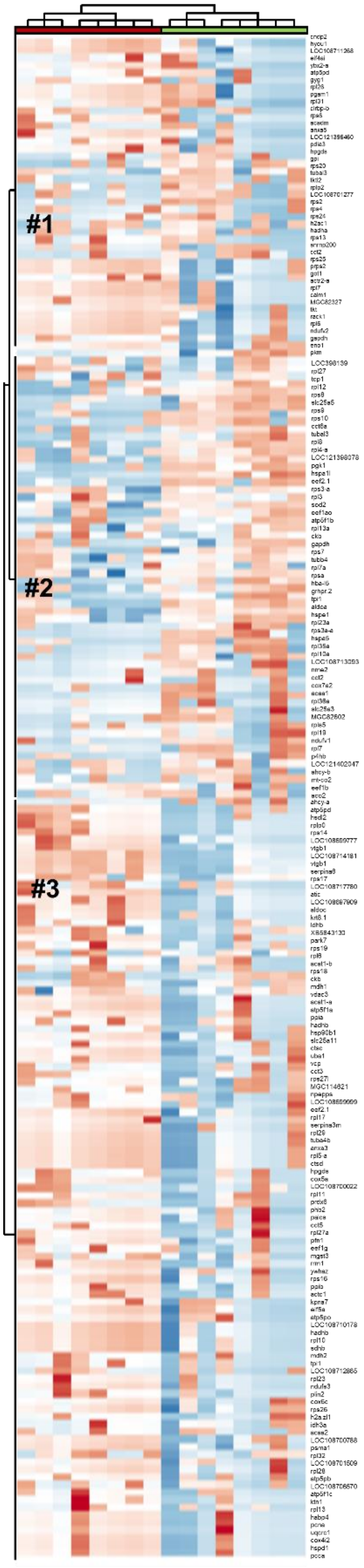

**Figure S7.** Close-up of the HCA-heat map of the top 200 most significantly differently differential proteins between the D11 and V11 clones at the stage-8 *X. laevis* blastula (**Fig. 5D**). The proteins are labeled with their corresponding gene names following the *Xenopus* naming nomenclature.
